# Structural basis of neurofibromin tetramerization and dimer-tetramer equilibrium

**DOI:** 10.1101/2025.02.13.638105

**Authors:** Shutian Si, Christian Tüting, Swanhild Lohse, Katharina Landfester, Ingo Lieberwirth, Panagiotis L. Kastritis, Anja Harder

**Affiliations:** Max-Planck Institute for Polymer Research, Mainz, Germany; Department of Integrative Structural Biochemistry, Institute of Biochemistry and Biotechnology, Martin Luther University Halle-Wittenberg, Halle/Saale 06120, Germany; Biozentrum and ZIK HALOmem, Martin Luther University Halle-Wittenberg, Halle/Saale 06120, Germany; CURE-NF Research Group, Medical Faculty, Martin Luther University Halle-Wittenberg, Halle (Saale), Germany; Institute of Chemical Biology, National Hellenic Research Foundation, Athens 11635, Greece; Medical Faculty, University of Muenster, Muenster, Germany

## Abstract

Human neurofibromin (NF1) is a tumor suppressor multidomain protein known to regulate cellular functions as a dimer. Using cryo-EM, we discovered that neurofibromin can form more highly organized structures and describe tetramerization and a dynamic dimer-to-tetramer equilibrium with extensive interlocked interfaces as well as structural flexibility. The new tetramer structure generates and modifies interaction surfaces and controls protein functions such as microtubule recruitment, thereby providing novel insights into cellular functions.

## Main

Neurofibromin is a tumor suppressor encoded by the *Neurofibromatosis type 1* (*NF1*) gene (OMIM *613113, 17q11.2). Pathogenic germline variants lead to the autosomal dominant tumor predisposition syndrome NF1 (OMIM 16220), additionally somatic variants have been detected in many different tumor types. Neurofibromin plays a fundamental role in development, signaling and homeostasis^1^. It consists of several domains including the GAP related domain (GRD) functioning as a Ras-GTPase activating protein (Ras-GAP)^2,3^. Still, detailed functional data are mainly available for the GRD and SecPH domains, leaving many protein functions unresolved. A better and precise dissection of the structural and functional mechanisms will improve developing molecular therapies for NF1-related malignancies as well as the numerous disease-associated complications. To date, cryo-electron microscopic analysis (cryo-EM) of neurofibromin resolved the highly intricate, multi-domain structure of its dimeric state^4-8^, organized on a 30 × 10 nm array of α-helices, forming a core α-solenoid scaffold with a figure-eight-like shape. The ∼640-kDa homodimer transitions between closed, autoinhibited conformations and open, active states that expose the Ras-binding site, as well as the SecPH domain for membrane sensing. This conformation change is regulated by nucleotide binding and interactions with the cellular membrane^6,7^. Pathogenic variants can disrupt stability, reduce expression or destabilize neurofibromin via dominant-negative effects, and severe effects in specific structural regions of the homodimer^8^ were shown to elucidate crucial protein functions^9^.

To deepen our understanding, the purified human recombinant wildtype neurofibromin isoform I of 2,818 aa (Uniprot ID#P21359-2 (UPI000002AEF8), missing 1371-1391 from the canonical sequence ID#P21359-1, isoform II) (**Methods, Fig. S1a**) was vitrified and visualized using cryo-EM by acquiring 24,960 micrographs in total (**Methods**). Clear 2D class averages of both the dimeric (**Fig. S1b)** and the unexpected, tetrameric form were visible (**Fig. 1a**). Statistical analysis of single particles shows the presence of one tetramer per four identified dimers (**Fig. S1c**), suggesting a specific equilibrium dynamic between dimer and tetramer formation. The 4:1 ratio implies that the affinity for tetramer formation is weaker relative to dimer formation and that its concentration linearly depends on dimer concentration. After extensive image processing (**Fig. S2**), the tetramer included 155,085 single particles, obeyed dihedral symmetry involving three two-fold axes, and was resolved to 4.26 Å resolution (FSC=0.143, **Table 1**). Local resolution spanned from 7.04 Å to 3.82 Å (**Fig. S3**), and, in some regions, individual amino acids were visible, especially after automated processing for additional noise reduction^10^ (**Fig. S4**).

**Table 1.** Cryo-EM data collection, refinement, and validation statistics.

|  | NF1 | NF1 lobe | NF1 core | NF1 tetramer | NF1 tetramer |
| --- | --- | --- | --- | --- | --- |
| **Data collection and processing** | 13,353 | 13,353 | 13,353 | 24,960 | 24,960 |
| Magnification | X105,000 | X105,000 | X105,000 | X105,000 | X105,000 |
| Voltage (kV) | 300 | 300 | 300 | 300 | 300 |
| Electron exposure (e-/Å<sup>2</sup>); DS denotes “dataset” | 40 (DS1)<br>49 (DS2) | 40 (DS1)<br>49 (DS2) | 40 (DS1)<br>49 (DS2) | 40 (DS1)<br>49 (DS2)<br>49 (DS3) | 40 (DS1)<br>49 (DS2)<br>49 (DS3) |
| Defocus range (μm) | -0.8 ~ -2.6 | -0.8 ~ -2.6 | -0.8 ~ -2.6 | -0.8 ~ -2.6 | -0.8 ~ -2.6 |
| Pixel size (Å) | 0.697 | 0.697 | 0.697 | 0.697<br>DS1&2)<br>1.394<br>(DS3) | 0.697<br>DS1&2)<br>1.394<br>(DS3) |
| Symmetry imposed | C1 | C1 | C1 | C1 | D2 |
| Initial particle images (no.) | 1,150,907 | 1,150,907 | 1,150,907 | 2,260,127 | 2,260,127 |
| Final particle images (no.) | 281,212 | 282,057 | 282,057 | 155,085 | 155,085 |
| Map resolution (Å)<br>FSC threshold | 3.35<br>(FSC=0.143) | 3.27<br>(FSC=0.143) | 3.26<br>(FSC=0.143) | 4.10<br>(FSC=0.143) | 4.26<br>(FSC=0.143) |
| Map resolution range (IQR, Å) | 2.96 – 3.68 | 2.81-3.49 | 2.95-3.60 | 3.59-4.65 | 3.82-7.04 |
| **Refinement** |  |  |  |  |  |
| Initial model used (PDB code) | 7R03 | 7R03 | 7R03 | 7R03 | 7R03 |
| Map sharpening *B* factor (Å<sup>2</sup>) | 0 (not modified) | 0 (not modified) | 0 (not modified) | 0 (not modified) | 0 (not modified) |
| Model composition |  |  |  |  |  |
| Non-hydrogen atoms | 35830 | 16992 | 8776 | 71660 | 71660 |
| Protein residues | 4514 | 2156 | 1132 | 9028 | 9028 |
| Ligands | 0 | 0 | 0 | 0 | 0 |
| *B* factors (Å<sup>2</sup>) |  |  |  |  |  |
| Protein | 89.6 | 91.1 | 73.8 | 87.2 | 139.2 |
| R.m.s. deviations |  |  |  |  |  |
| Bond lengths (Å) | 0.005 | 0.004 | 0.003 | 0.003 | 0.003 |
| Bond angles (°) | 0.856 | 0.781 | 0.763 | 0.664 | 0.730 |
| Validation |  |  |  |  |  |
| MolProbity score | 2.11 | 1.96 | 1.89 | 1.89 | 2.01 |
| Clashscore | 18.81 | 13.91 | 14.16 | 12.13 | 15.57 |
| Rotamer outliers (%) | 0.70 | 0.89 | 1.00 | 0.01 | 0.04 |
| Poor rotamers (%) |  |  |  |  |  |
| Ramachandran plot |  |  |  |  |  |
| Favored (%) | 95.07 | 95.51 | 96.51 | 95.81 | 95.53 |
| Allowed (%) | 4.84 | 4.44 | 3.49 | 4.10 | 4.36 |
| Disallowed (%) | 0.09 | 0.05 | 0.00 | 0.09 | 0.11 |

**Figure 1.**
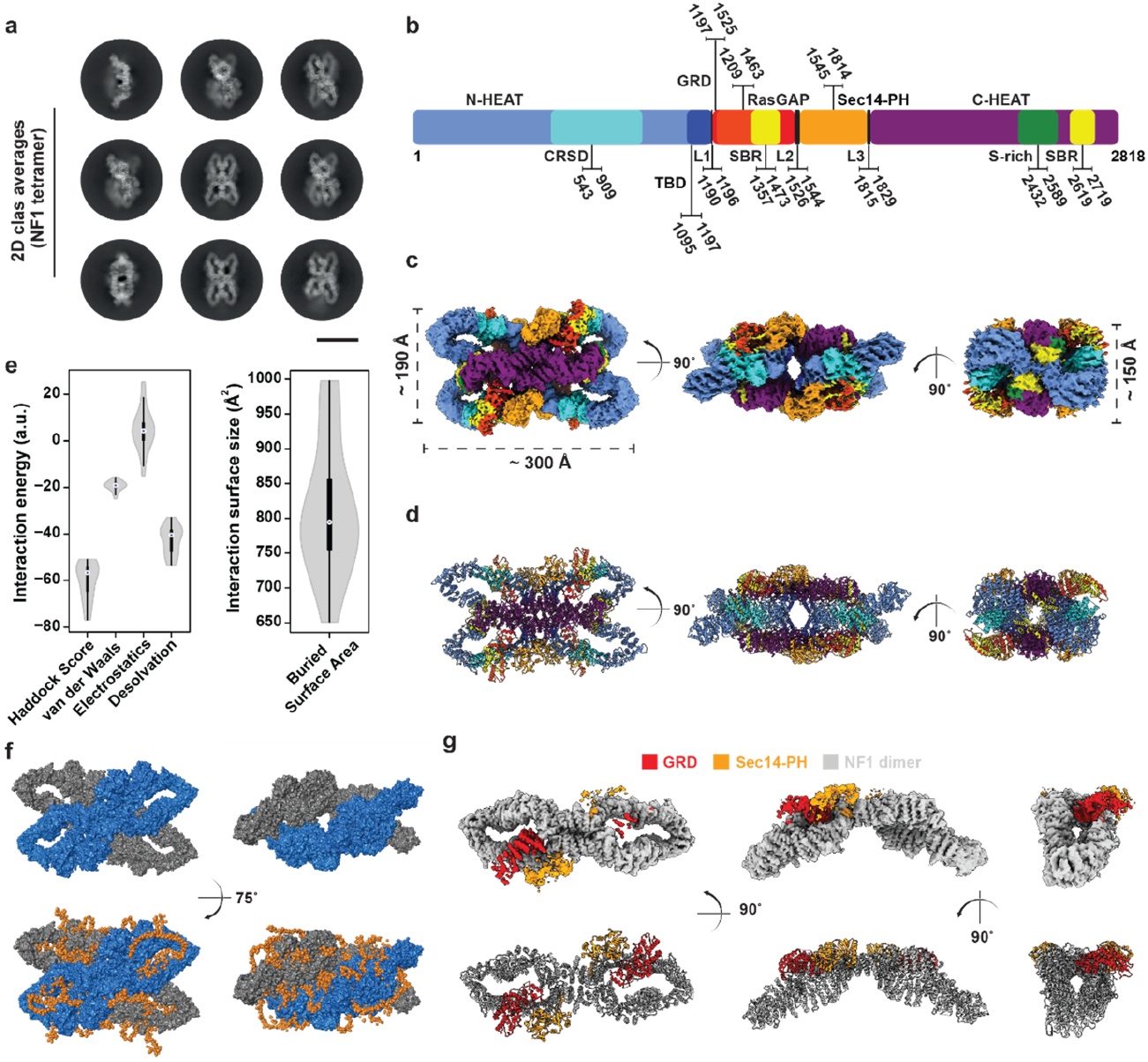
Characterization of the NF1 tetramer structure. **a**, Representative 2D classes of NF1 tetramer are shown. The scale bar indicates 20nm. **b**, The scheme illustrates the domain organization of NF-1. **c**, The 4.26Å cryo-EM map of NF1 tetramer in three different views, the color of domain corresponds to residue ranges shown in the domain scheme, **b**. **d**, Ribbon model of the NF1 tetramer. **e**, Energetic contributions for 20 generated refined models utilizing HADDOCK. On the left, violin plots for HADDOCK score, van der Waals, Electrostatics, and Desolvation energies are shown; on the right, a violin plot is shown with the calculated values of the buried surface area in Å^2^. **f**, On top, the resolved NF1 tetramer structure is shown as surface representation. Each neurofibromin dimer is shown in blue and grey colors. On the bottom, low confidence regions (<30 pLDTT score) of the superimposed Alphafold 3 model of the monomer are shown with orange. These loop regions are shown as sphere representation and cover the observed cavities in the determined complex. **g**, The resolved dimeric neurofibromin in different views. The GRD and Sec14-PH regions are colored in red and orange separately.

The multi-domain neurofibromin monomer (**Fig. 1b**) molds a flexible D2 homotetramer (**Fig. 1c**) formed by the association of two C2 homodimers (**Fig. 1c,d**) with an exposed surface of an impressive ∼400000 Å^2^. Each homodimer exhibits twofold rotational symmetry with associated flexibility and known conformation changes^5-8^. Molecular modelling with available structural data (**Methods**) and AlphaFold3^11^, and subsequent map refinement of both resolved dimer (3.35 Å, FSC=0.143, **Table 1**) and tetramer (**Methods, Table 1**), revealed an extensive inter-dimer interface of 6081.1 Å^2^. This extended interface is characterized by complementary surfaces, leading to a tightly interlocked structure of two lemniscates (**Fig. 1d**). The opposing dimer surfaces fit together by maximizing the interface size, forming a cohesive tetrameric assembly.

A prominent feature of this large assembly is its relatively small buried surface area (BSA, 813 ± 96 Å^2^) (**Fig. 1e**), *i*.*e*., the surface of residues that directly interact in the bound state, which explains low binding affinities^12^. The complex is governed mostly by desolvation energy (**Fig. 1e**), implying that removal of water for interface formation is more beneficial for binding as compared to, e.g., electrostatics or van der Waals, that contribute less to complexation (**Fig. 1e**). Overall, such energetic contributions qualitatively describe a complex that binds in the high μm range^13^, in agreement with the observed abundance of the dimeric form as compared to the NF1 tetramer (**Fig. S1c**). The higher-order structure also forms a central pore (**Fig. 1d, f, Fig. S5, Fig. S6**), which is organized by the C-ter of the solenoid and flexible loops (**Fig. S5, Fig. S6**). The AlphaFold3 monomeric model, when superimposed on the resolved structure, shows that the formed pores of the neurofibromin tetramer are accommodated by flexible linkers (**Fig. 1f**). The inferred presence of disorder suggests that extensive, low-complexity interactions further contribute to the tetramer stability.

The heterogeneity of classified particles (**Fig. 1a, Fig S1b**), and local resolution calculations of final C1 and D2-symmetrized maps **(Fig. S3)** indicated substantial variability to exist between the interfaces. To address this, 3D variability analysis^14^ was used for the symmetrized map to generate a consecutive volume series of 20 frames. Results show the presence of gradual rigid-body displacement of neurofibromin dimers, specifically following shear-based/scissor-like conformational change (**Movie S1**). Shearing motions are well-described mechanisms for multimeric proteins that incorporate allosteric communication across subunits, are prominent in proteins composed by repeat elements^15^, and especially common in α-solenoids^16^. Neurofibromin tetramer core motions involve HEAT repeat α-helices (E1034-M1050, L1064-L1080, E1094-E1119 and S1135-N1156) which displace by ∼10 Å during shearing (**Movie S2**). Lobes of the tetramer are less resolved (**Movie S1, Fig. S3**), indicating that described flexibility of the dimer-embedded NF1 HEAT repeats^5-8^ are retained in the higher-order tetrameric structure. HEAT repeats frequently induce such “sliding” effects in other large biomolecular complexes that involve a high HEAT repeat content^17,18^, similar to neurofibromin.

To assess the extent of conformational changes neurofibromin might undergo from the unbound to the bound state, we also resolved the more abundant dimer in the sample (**Table 1**). Resolution for the whole dimer reached 3.35 Å (FSC=0.143), with local refinement and reconstruction of the core and lobe regions reaching 3.26 Å and 3.27 Å (FSC=0.143), respectively (**Table 1, Fig. S2, Fig. S3**). Cross-molecule superpositions of resolved models showed minimal conformation changes (RMSD < 1 Å, **Fig. S7**), therefore, pointing to a lock-and-key mechanism for tetramer formation with potentially minor side-chain re-adjustments. Overall, both GRD and SecPH modules could be fitted unequivocally into either dimer or tetrameric densities. Therefore, resolved conformation was very similar to the auto-inhibited neurofibromin dimer state with occluded Ras-binding sites^7^ (**Fig. 1g**). However, due to the presence of 4 GRDs in the tetramer, and the extensive conformational flexibility of the higher-order structure (**Movie S1**), we cannot exclude the presence of other conformations. The positioning of the proximal SecPH module, responsible for lipid binding and membrane recruitment, is also in the closed, auto-inhibited conformation, as expected (**Fig. 1g**).

To understand the consequences of the tetramer for protein-protein interactions involving neurofibromin, Alphafold 3-based systematic modelling of all major protein-protein interfaces (PPIs) (*N*=52), recently reviewed^19^, was performed (**Methods**). Filtering of the interaction models according to a conservative threshold of predicted aligned error (PAE) score (<20 a.u.) and proximity calculations of residues (d_Cα_-_Cα_ <10 Å) predicts potential interfaces for 35 of those (56%, **Fig. 2a**). Overall, 598 residues localize in extended sub-surface patches across neurofibromin and are involved in the recruitment of those 35 interactors (**Fig. 2b)**. Importantly, substantial portion of those patches is buried upon tetramerization (**Fig. 2c-d**), revealing obvious functional differences for recruitment of interactors. A prominent example is recruitment of neurofibromin on microtubules^20-22^, which impacts RAS-GAP activity^22^. The neurofibromin dimer adopts an extended, curved conformation, interacting with the external surface of the microtubule, with the tubulin-binding domain (TBD) at the interface core, while exposing the GRD and SecPH modules. It aligns along the protofilament axis, making multiple contact points with tubulin subunits through its HEAT repeats (**Fig. 2c**). In contrast, the tetrameric form cannot bind, as the additional dimeric α-solenoid fold extensively clashes with the microtubule surface, preventing association (**Fig. 2d**). Buried sites are also predicted for estrogen receptor, 14-3-3 proteins, and syndecans, suggesting selected functions depending on structure in cell cycle regulation or in specific cell types, e.g. neurons or hormone sensitive cells.

**Figure 2.**
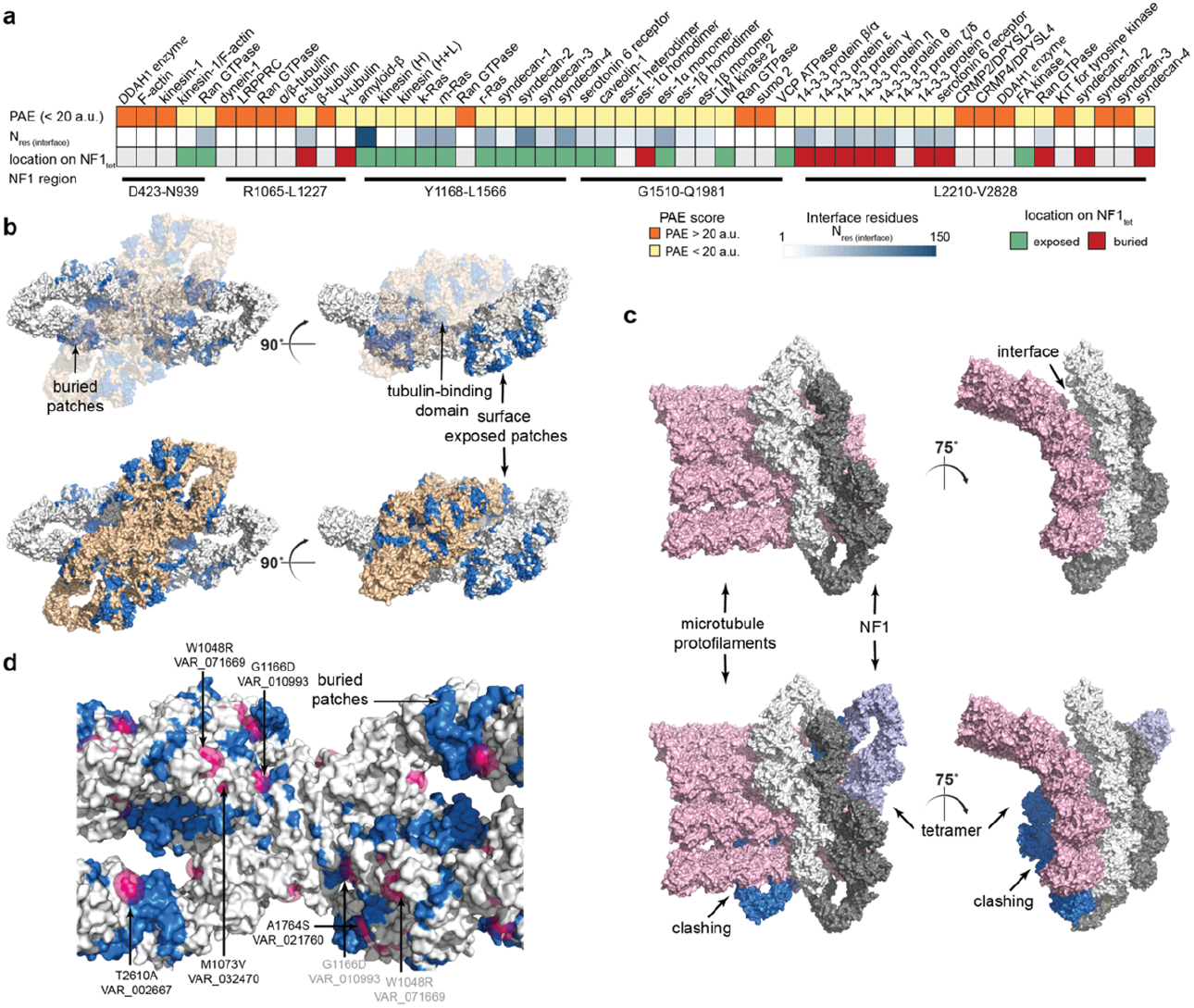
Implications of NF1 tetramerization in protein-protein interactions and localization of SNPs. **a**, Heatmap showing the confidence and resulting interface assessment for predicted binders of NF1. **b**, Mapping of interface areas on top of the NF1 tetramer, highlighting exposed and buried regions; The tubulin-binding domain is also shown, all color-coded with blue. **c**, A binding model of NF1 with microtubules, based on the predicted NF1 domain – tubulin interface; It is evident that upon tetramerization, major clashes are present in the structure, while the dimeric NF1 can be accommodated on microtubules with extensive surface complementarity. **d**, Mapping of SNPs (purple color) retrieved from Uniprot on top of the NF1 tetrameric interface. Regions of protein-protein interactions with confident PAE scores (< 20 a.u.) are shown in blue. Models shown are rendered in PyMOL v2.5 with surface representation.

Functional differences are also evident when mapping a selection of carefully curated *NF1* single nucleotide polymorphisms (SNPs), present in Uniprot (P21359), when the dimeric and tetrameric form are considered. Out of 87 variants, few (*N*=8, H31R, G629R, S665F, F1193C, L1196R, S1468G, Y2171N, T2486I) are localized in unresolved regions, while the vast majority (*N*=71) are surface exposed in both dimer and tetramer states. This shows that the exposed tetramer surface is more prone to variation as opposed to the tetrameric interface, a general attribute of biological interfaces^23^. Interestingly, the remaining missense variants localize at, or in proximity to, the tetrameric interface (*N*=8, W1048R, M1073V, L1147P, N1156S, G1166D,

A1764S, D1991N, T2610A). These variants are exposed in the dimeric state, but buried in the tetrameric state, e.g. W1048R, M1073V, G1166D, A1764S and T2610A (**Fig. 2d**). Such a differential localization within in the structure might impact dimer-to-tetramer equilibria, possibly existing *in vivo* with functional consequences for binder recruitment, and therefore, the multi-faceted role of neurofibromin. For example, variants T2610A, M1073V, and N1156S, although of conflicting pathogenicities, have been described in specific phenotypes including pulmonary stenosis, juvenile myelomonocytic leukemia, and NF-Noonan- or cardiovascular phenotypes of NF1.

To conclude, the counts of the single particles showed an approximate 4:1 presence (dimers:tetramers) indicating a simple equilibrium *in vitro*. In addition, the minor conformational changes that neurofibromin undergoes upon tetrameric assembly suggests government mostly by enthalpic, rather than entropic forces^24^. Its buried surface area, 813 ± 96 Å^2^, if translated into binding free energy, shows a loose binding of the two dimers (at the μM range^13^); however, the inherent flexibility of the structure, as shown by variability analysis, may affect complexation by thermal motion, which would induce allostery via entropic, long-range effects^25^, which would increase off rates of the tetramer^24^. Such relative populations of dimer-to-tetramer equilibria were observed in other biological molecules, with the most prominent example being hemoglobin^26^. This parallelism underscores the evolutionary utility of dimer-to-tetramer equilibria in regulating protein function through higher-order multimerization. Distinct binders or local concentration effects can simply regulate the equilibrium, as well as possible physical-chemical conditions, such as low temperatures, stress, or local pH changes, with in-cell implications, *e*.*g*., recruitment on microtubules.

To summarize, we demonstrate tetramerization of neurofibromin, pointing to functional regulation via multimerization. This discovery of a new higher organized interlocked structure widens our understanding of importance of this protein and points to novel neurofibromin functions.

## Online Methods

### Protein expression and purification

The full-length recombinant wildtype human isoform I of neurofibromin (2,828 aa; Uniprot ID#P21359-2) was expressed and purified in a baculovirus-insect cell system tagged with N-His/Strep, and using a TEV protease cleavage site by CREATIVE BIOMART® (Shirley, NY 11967, USA). The purified protein (in 50 mM Tris, pH 8.0, 200mM NaCl, 2mM MgCl2, 2mM DTT) was stored at -80°C and aliquoted for further use.

For purification, cells were resuspended in 100 ml lysis buffer and broke by ultrasound (1s on, 1s off, 30% power, for 20 minutes), and centrifuged thereafter at 11900 rpm for 20 minutes. The supernatant with incubated with pre-equilibrated Ni-NTA affinity chromatography was column, washed twice with Binding Buffer (50 mM Tris, 200 mM NaCl, 20 mM Imidazole, pH 8.0 (Lot 20240726), eluted using Ni-NTA Elution Buffer (50 mM Tris, 200 mM NaCl, 300 mM Imidazole, pH 8.0 (Lot 20240726)), and collected. SDS-PAGE analysis revealed requirement of re-purification as described. A third purification step was added using SEC200-24ml purification. The concentrated supernatant was taken from the gel and loaded onto the pre-equilibrated SEC200-24ml column with Buffer (50 mM Tris pH 8.0, 200 mM NaCl, 2 mM MgCl2, 2 mM DTT). One ml using a loop at a flow rate of 0.4 ml/min was collected to perform SDS-PAGE analysis.

For protein analysis, neurofibromin was dialyzed and concentrated, followed by Bradford, SDS-PAGE, Western Blot and HPLC detection. The protein of 1.04mg/ml (by Bradford) was run by 4-20% Coomassie Blue stained SDS-PAGE and showed > 85% purity. For verification of the calculated 320 kDa protein SDS-PAGE was performed using an Anti-His Mouse Monoclonal Antibody due to manufacturers protocols (**Fig. S1a**).

### Cryo-EM specimen preparation and data collection

For cryo-TEM examination the samples were vitrified using a Vitrobot Mark IV (Thermo Fisher, Hilsboro Oregon) plunging device at 4°C and 100% humidity. The protein was diluted to 0.5mg/ml in a buffer containing 50mM Tris, pH 8.0, 200mM NaCl, 2mM MgCl2, 2mM DTT. 3 µl of the protein sample was applied to Quantifoil grid (Au R1.2/1.3 300 mesh and Cu R2/1 300 mesh) that was glow discharged in an oxygen plasma cleaner (Diener Nano®, Diener electronic, Germany) shortly before. After removing excess sample solution with a blot force of -1 for 2.5s, the grid is immediately plunged into liquid ethane. For the subsequent examination the specimen is transferred to a TEM (FEI Titan Krios G4) keeping cryogenic conditions.

Data collection was done using an acceleration voltage of 300 kV. Micrographs were acquired with a 4k Direct Electron Detection Camera (Gatan K3) under low dose conditions. A normal magnification of X105,000 was used for the data recording, corresponding to pixel size of 0.697Å for two datasets and 1.394Å for the third dataset (bin 2). Defocus values varied from -0.8 to -2.6 μm during acquisition. For the first data set, 41-frame movies were collected in super-resolution mode with 3.28s of total exposure, leading to a total electron dose of 40 e-/Å^2^. For the second data set, 42-frame movies were collected in super-resolution mode with 3.80s of total exposure, leading to a total electron dose of 49 e-/Å^2^. For the third data set, 42-frame movies were collected in super-resolution mode with 3.75s of total exposure, leading to a total electron dose of 49 e-/Å^2^. Extended Data Table 1 summarizes the model statistics.

### Cryo-EM data processing of neurofibromin dimer and tetramer

Two data sets were collected and accessed for NF1 dimer, corresponding to 13,353 micrographs. The micrographs were x2 binned, generating a pixel size of 1.394 Å per pixel. The data was loaded into CryoSPARC^27^ to calculate motion correction and contrast transfer function (CTF) estimation. A total of 5,505,757 particles were automatically picked using template picking. Meanwhile the micrographs were carefully screened to remove low-quality data. The particles were extracted in a box of 400 pixels and crop to box size of 100 pixels. For each data set, after several rounds of two-dimensional (2D) classification and heterogeneous refinement, the selected particles and volume were subjected to homogeneous refinement and reference motion correction. 161,982 particles from data set 1 and 300,810 particles from data set 2 were combined together for further Heterogeneous refinement. Finally, 281,212 good particles were subjected to homogeneous refinement. The final improved reconstruction map was refined to 3.39 Å (FSC=0.143). Further local refinement led to a final dimer reconstruction at 3.35 Å resolution with C1 symmetry. For reconstruction of the lobe region and core region, particles derived from the reference motion correction were subjected to heterogeneous refinements and manually particle removal. The volume and mask for lobe region were done with Chimera. 312,775 particles were subjected to signal subtraction, followed by further CTF refinement and manually lower quality particle removal. Finally, 282,057 particles yielded a 3.27 Å overall resolution. With the similar procedure, 282,057 particles yielded a 3.26 Å overall resolution for core region. To improve the interpretability of the Cryo-EM map, the final reconstructions for NF1 homodimer were sharpened using DeepEMhancer^10^.

For NF1 tetramer, extra third data set in which 11,607 micrographs with a pixel size of 1.394 Å were collected and combined with the previous two data sets. Low quality micrographs were then manually removed. 545,463 particles of NF1 tetramer were automatically picked using template picking and subjected to multiple rounds of 2D classification and heterogeneous refinement. Finally, 155,085 particles were subjected to homogeneous refinement with or without D2 symmetry. The resolution for final maps reached to 4.10 Å and 4.26 Å, after applying C1 and D2 symmetry respectively. 3D variability analysis^14^ was performed by default, defining 20 different classes to be calculated

### Model building and refinement

The published model of NF1^7^ (PDB ID: 7R03), having identical sequence to our designed NF1 construct, was rigid-body fitted in the best resolved density map (lobe region) using UCSF Chimera^28^ and manually adjusted in Coot^29^, during which several residues of linker regions were added based on the local density. The model was further subjected to several rounds of real-space refinements in Phenix1.20.1-4487^30^ using maps generated by DeepEMhancer^10^ and manual refinements. Similarly, the NF1 whole length and core region were manually adjusted in Coot using the previously obtained model and real-space-refined in Phenix using corresponding maps generated by DeepEMhancer^10^.

### Protein modelling and energy calculations

Energy calculations were performed using the HADDOCK webserver^31^, version 2.4^32^, utilizing the refinement interface by default. Analysis of all 20 generated and refined tetrameric models was performed to plot distributions of non-covalent interactions and corresponding buried surface areas. Surface area calculations were performed in PyMOL v2.5 (https://www.pymol.org/) using the *get_area* function for the unbound dimers and the bound tetramer, with calculations as previously described^33^. To map highly flexible regions on top of the structural model derived from cryo-EM, the full NF1 sequence was used to be modelled by AlphaFold3^11^; Then, superposition of the derived model with the atomic model of the NF1 tetramer was performed in PyMOL v2.5.

To model all potential protein-protein interaction of NF1, modelling with AlphaFold3^11^ was performed. Briefly, the corresponding sequence regions of NF1 were retrieved from the literature^19^. Then, extracted NF1 domains and the corresponding full Uniprot^34^ sequences of reported binders were used for modelling. To extract accurate interface residues, PAE scores lower than 20 *a*.*u*. were considered, and proximity of residues (d_Cα_-_Cα_ < 10 Å) was used to filter the remaining interactions. For modelling the microtubule-NF1 complex, AlphaFold prediction of tubulin were superimposed to the atomic model of the resolved microtubule structure^35^ (PDB ID: 3J6E). For the SNP mapping, the data were retrieved from Uniprot, aligned to match the sequence of the NF1 isoform 2, and visualized in PyMOL v2.5 to assess their surface accessibility in the dimeric and tetrameric states.

## Data availability

The maps will be available in the Electron Microscopy Data Bank (EMDB). The atomic models will be available in the Protein Data Bank database. Original movies and picked particles will be available in EMPIAR.

## Acknowledgements

This work was supported by the European Union through funding of the Horizon Europe ERA Chair “hot4cryo” project number 101086665 (to PLK), the Federal Ministry for Education and Research (BMBF, ZIK program; Grant nos. 03Z22HN23 and 03COV04 to PLK), the European Regional Development Funds for Saxony-Anhalt (grant nos. EFRE: ZS/2016/04/78115 and ZS/2024/05/187255; and to PLK), the Federal State Saxony-Anhalt (to PLK), funding by DFG (project number 391498659, RTG 2467 to PLK), the region of Saxony-Anhalt, and the Martin-Luther University of Halle-Wittenberg (to PLK). Additionally, this work was supported by the Bundesverband Neurofibromatosen e.V. (NF1466), Nothing-is-forever e.V. (NiF489) and private sponsorings (CureNF) to AH (project numbers 33103058, 33103052, 30101018). SS would like to acknowledge funding from the CRC1551 “Polymer concepts in cellular function” of the Deutsche Forschungsgemeinschaft (project number 464588647). IL and KL would like to thank the Max-Planck Society for financial support. We thank Olha Storozhuk for preparatory work.

## Author information

These authors contributed equally: Ingo Lieberwirth, Panagiotis L. Kastritis, and Anja Harder.

## Contributions

AH organized and facilitated the implementation of the study. SS performed the cryo-EM experiments, acquired, processed and analyzed the data. CT aided in data processing. SS, CT, and PK carried out modeling, map interpretation and performed integrative structure calculations. CT performed Alphafold modelling. SL performed analysis of structural data and contributed to the manuscript. KL and IL supervised the cryo-EM investigations and provided the imaging and analysis platform. PLK, SS, and AH drafted the manuscript with input from all co-authors. AH, IL and PLK designed and supervised the study in detail and acquired funding. All authors approved the draft.

## Ethics Declaration

The authors declare no competing interests. Ethics approval has been granted by the ethics committee of the Medical Faculty at Martin-Luther-University (2021-177) and the Rhineland-Palatinate State Medical Centre (2024-17491).

