## Supplementary Figures and Movies for "Structural basis of neurofibromin tetramerization and dimer-tetramer equilibrium"

| | |
|---|---|
| Shutian Si | |
| Christian Tüting | |
| Swanhild Lohse | |
| Katharina Landfester | |
| Ingo Lieberwirth | |
| Panagiotis L. Kastritis | |
| Anja Harder | |

* Corresponding authors

Correspondence should be addressed to either of these authors:

**Ingo Lieberwirth** – Max-Planck Institute for Polymer Research, Ackermannweg 10, 55128, Mainz, Germany,,

**Panagiotis L. Kastritis** – Martin Luther University Halle-Wittenberg, Institute of Biochemistry and Biotechnology, Department of Integrative Structural Biochemistry, Weinbergweg 20, 06120 Halle (Saale), Germany,,

**Anja Harder** – Martin Luther University Halle-Wittenberg, Medical Faculty, Institute of Anatomy and Cell Biology, CURE-NF Research Group, Große Steinstrasse 52, 06112 Halle (Saale), Germany,.

### Supplementary Figures

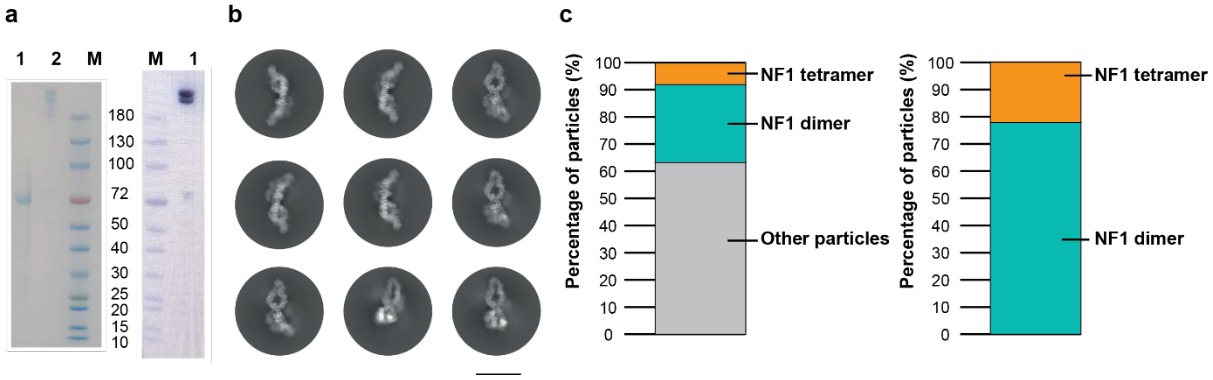


**Supplementary Figure 1: Purification, characterization, and statistical analysis of single particles for human recombinant neurofibromin.**
**a**, SDS page and Western blot for purification and size of human recombinant wildtype (wt) neurofibromin (NF1). Indicated are lane 1 - BSA, line 2 – wt NF1, M - protein molecular weight marker (10, 15, 20, 25, 30, 40, 50, 72, 100, 150, and 180 kD.) **b**, 2D average classes of NF1 dimer. The scale bar indicates 20 nm. **c**, Statistical analysis of single particles using CryoSPARC v4.6.0 shows the ratio between different particle classes.

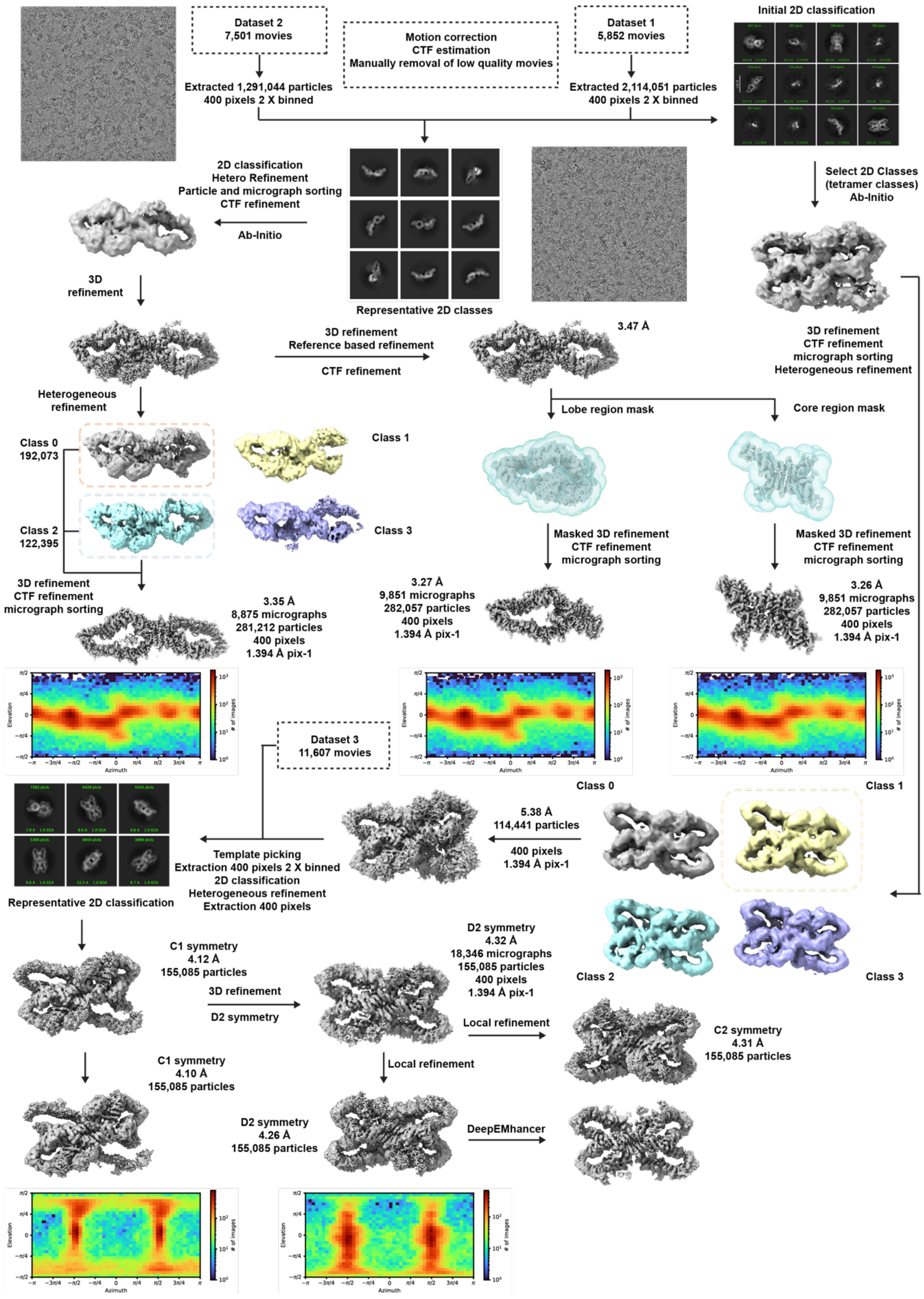


**Supplementary Figure 2: cryo-EM image processing workflow.**

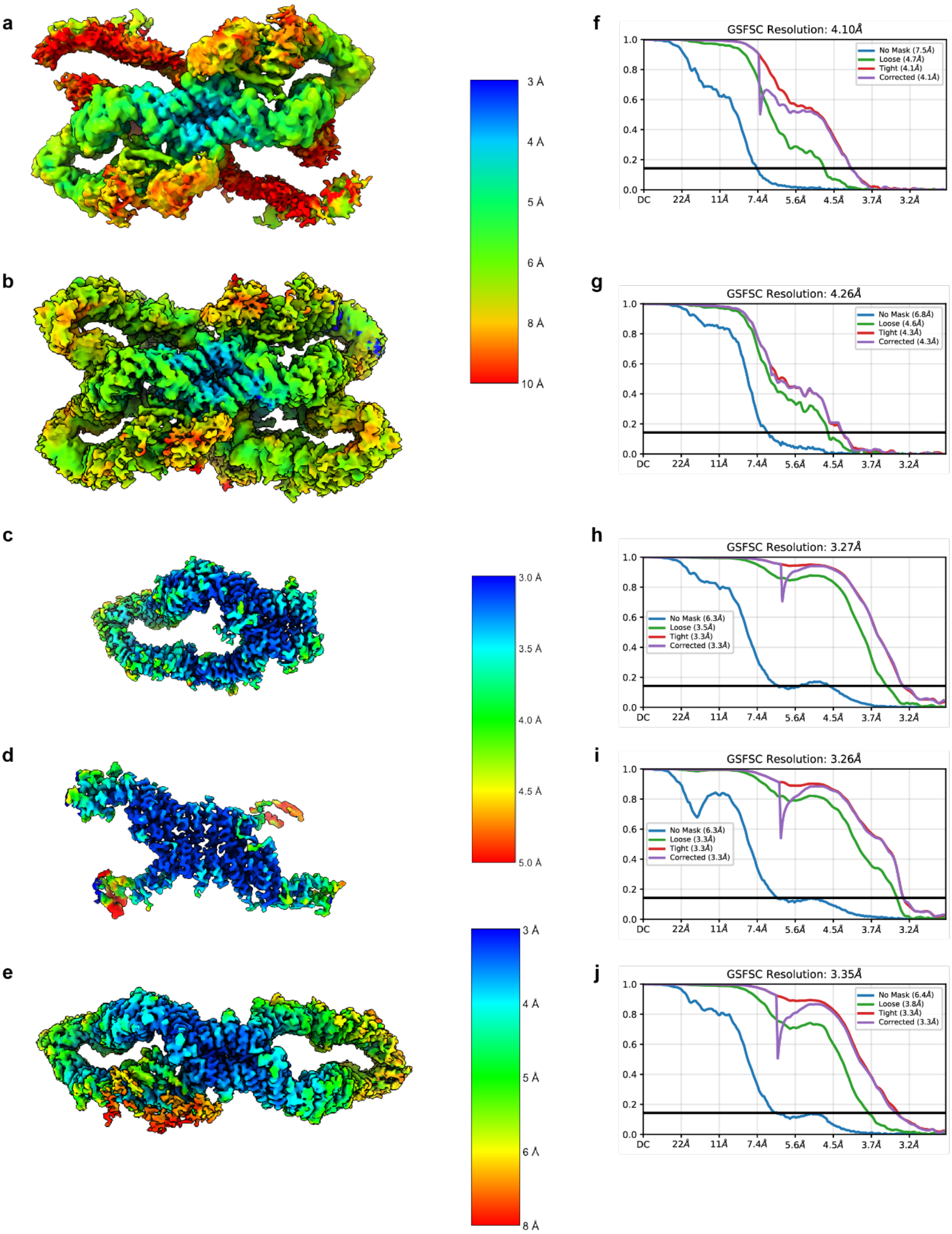


**Supplementary Figure 3: Local resolution of final reconstructions.**
Final 3D reconstructions colored based on local resolution estimation calculated by cryoSPARC. Local resolution estimation for **a**, NF1 tetramer, **b**, NF1 tetramer with D2 symmetry, **c**, NF1 lobe region, **d**, NF1 core region, **e**, NF1 dimer. **f-j**, The estimated resolution (FSC=0.143) plot for each solved reconstruction.

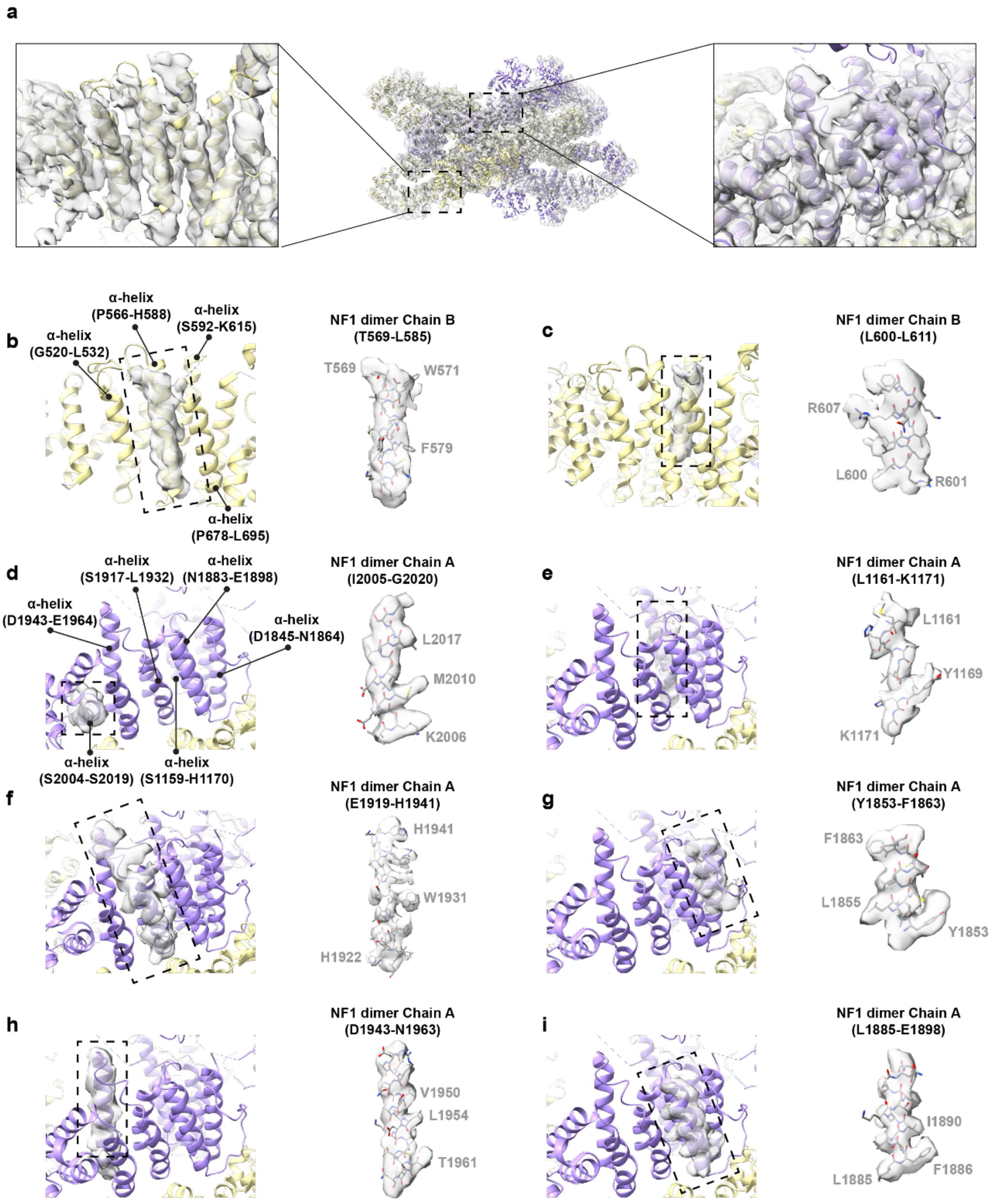


**Supplementary Figure 4: Quality of cryo-EM density maps.**
**a**, Overlay of structure model on cryo-EM density map for NF1 tetramer. The 4.26 Å reconstruction of NF1 tetramer with D2 symmetry was processed by DeepEMhancer. Two NF1 dimers are marked by purple and yellow respectively. Zoom-in views show the lobe region in the left box and central core region in the right box. **b**,**c** Representative densities of side chains for lobe region, and **d**-**i** for the core region.

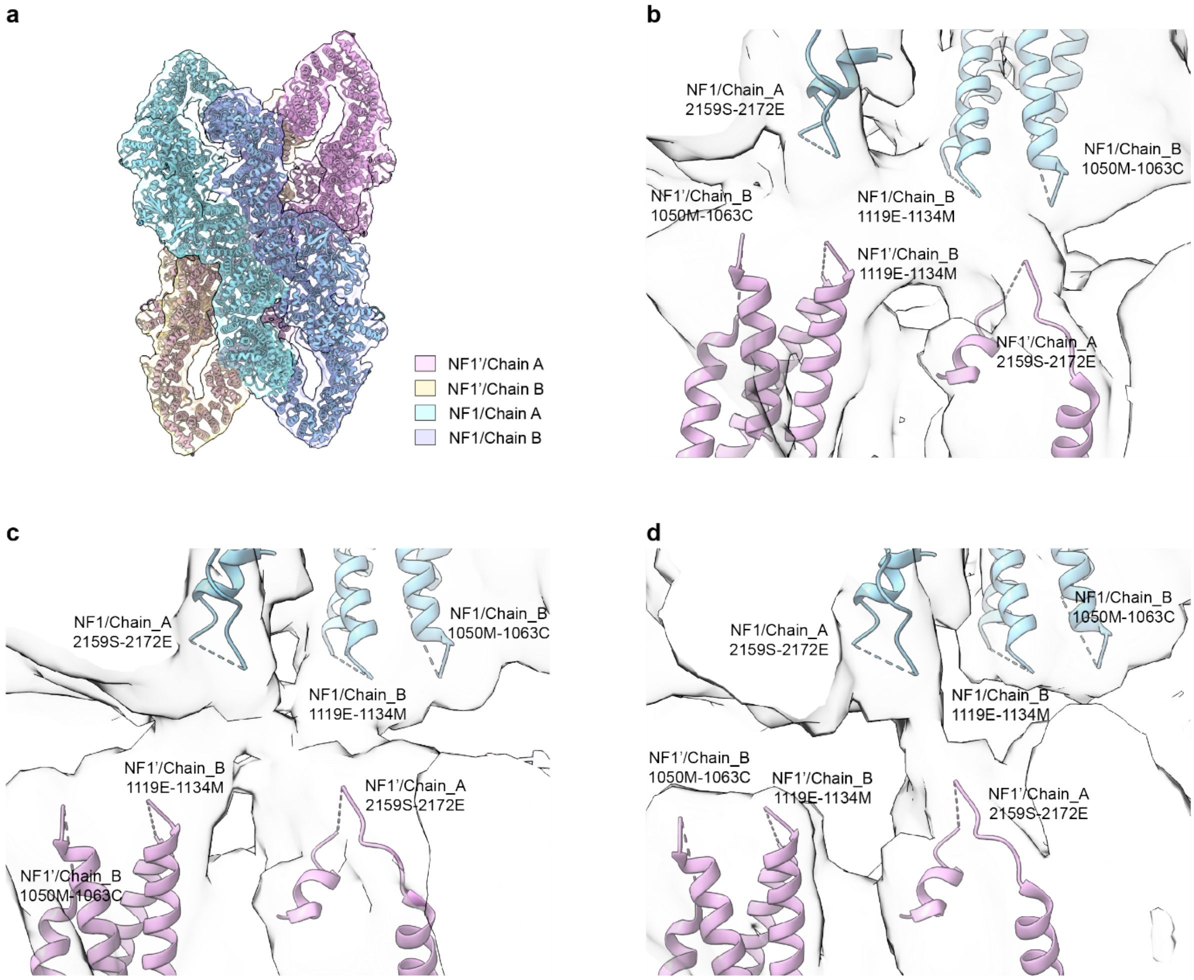


**Supplementary Figure 5: Core region dynamics of NF1 tetramer when fit model into 3D volume series frames.**

3D variability analysis shows the dynamics of NF1 tetramer core region when fitting model into different volume series frames. The missing loops were labeled with dashed lines and corresponding sequences. **a**, Labelling of the side chains of NF1 tetramer with different colors; **b**, NF1 homodimer fitting into 3D Var volume frame 000, the middle volume frame 010, **c**, and volume frame 019, **d**.

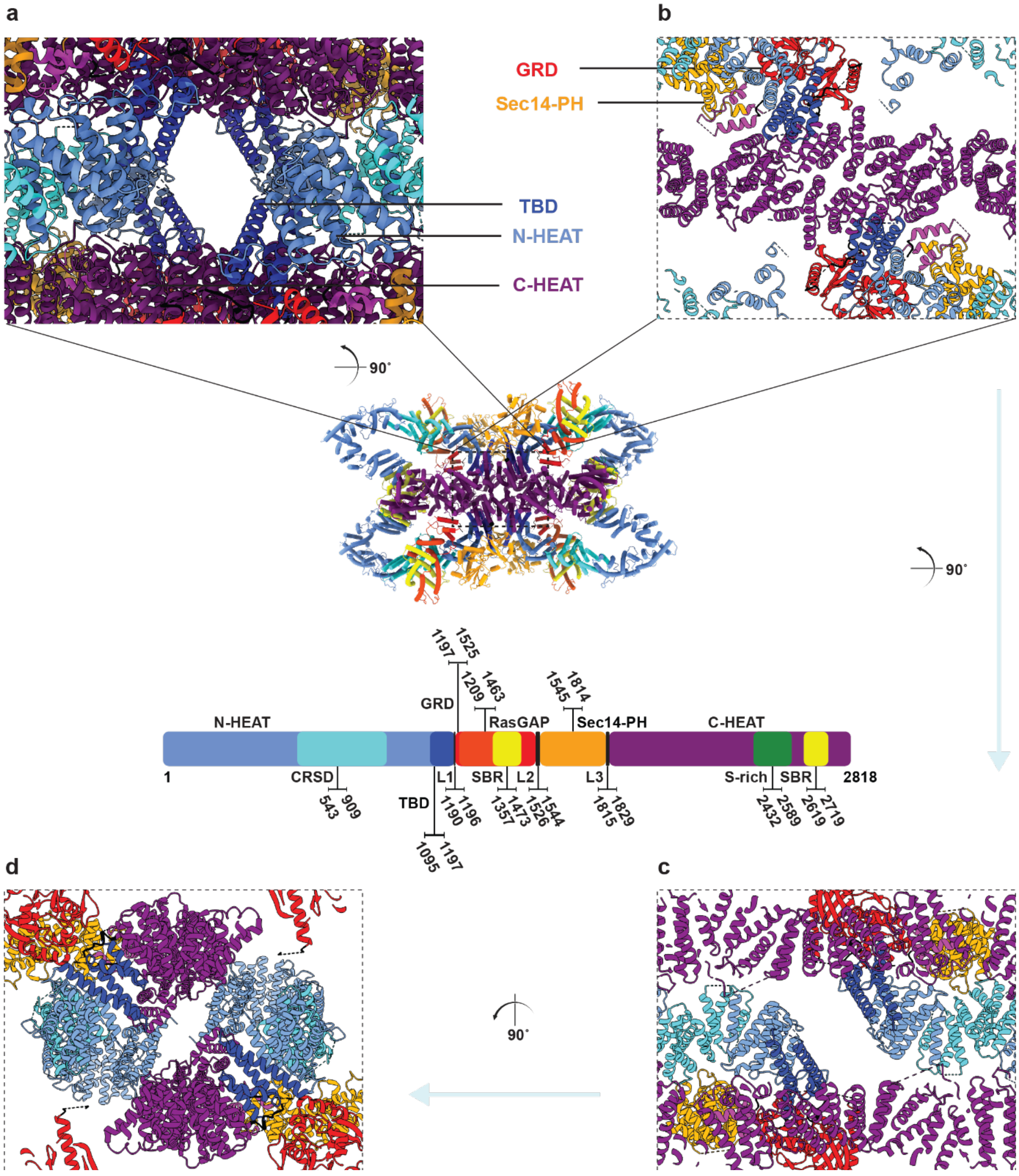


**Supplementary Figure 6: Overview of the central cavity.**
Formation of central cavity in the D2 symmetry axis. **a**, Side view of the central cavity. **b**-**d**, Cross-section views in three different dimensions.

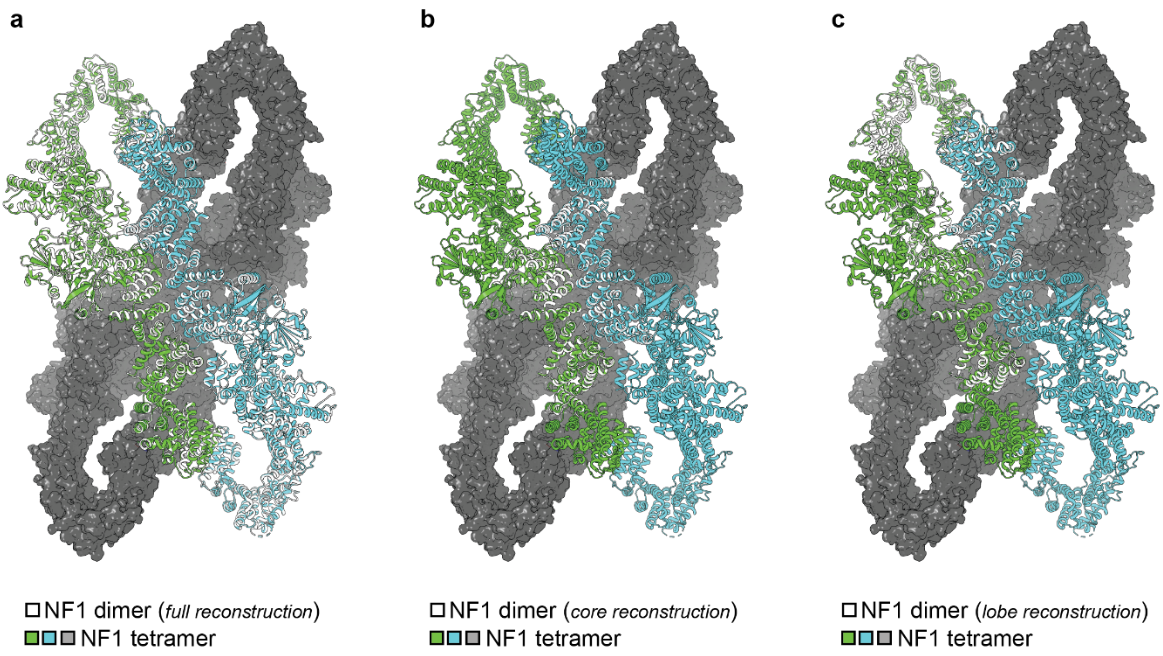


**Supplementary Figure 7: Cross-molecule superpositions of resolved models.**
The NF1 monomers within the NF1 tetramer complex are colored in green and blue. **a**, Superimposition of NF1 dimer (white) onto NF1 tetramer. **b**, Superimposition of core region (white) of NF1 dimer onto NF1 tetramer. **c**, Superimposition of lobe region (white) of NF1 dimer onto NF1 tetramer.

### Supplementary Videos (Movies)

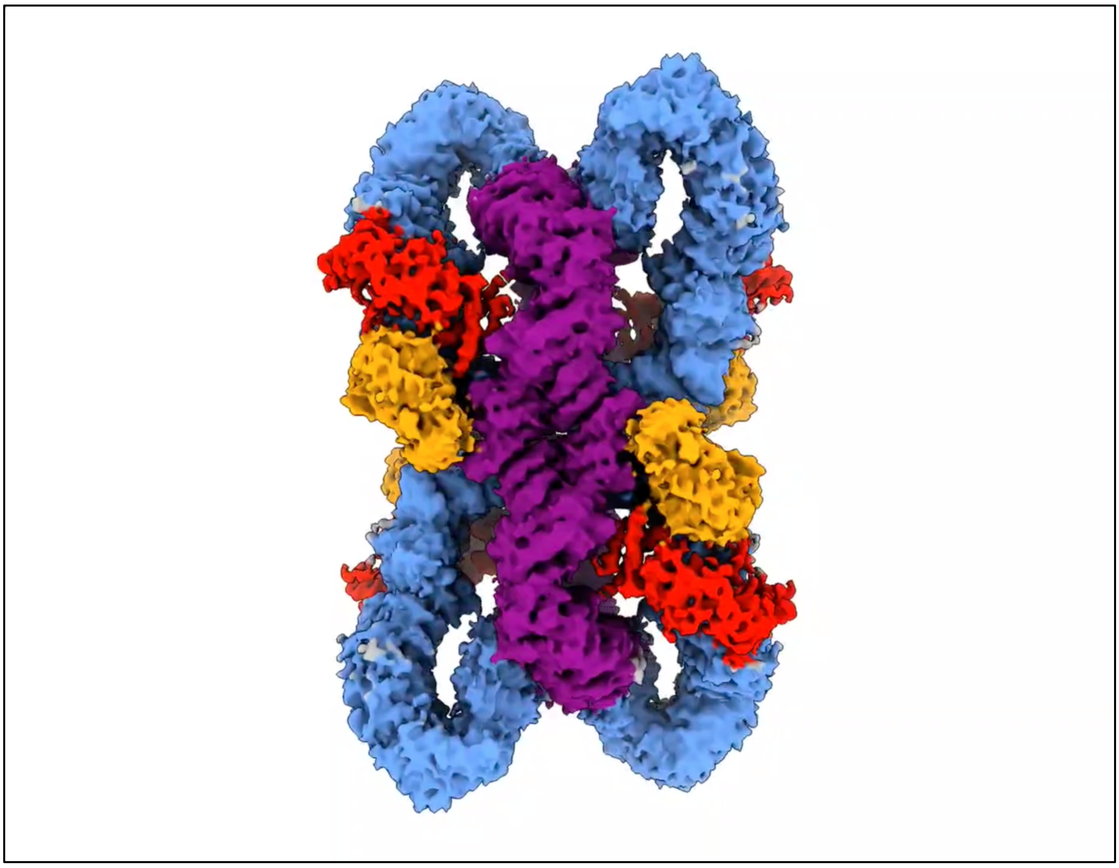

**Movie S1.** 3D variability analysis of the NF1 tetramer, along with local resolution estimation and domain annotation. Link to download: https://keeper.mpdl.mpg.de/f/7d796c32e7b14dceb6ff/?dl=1

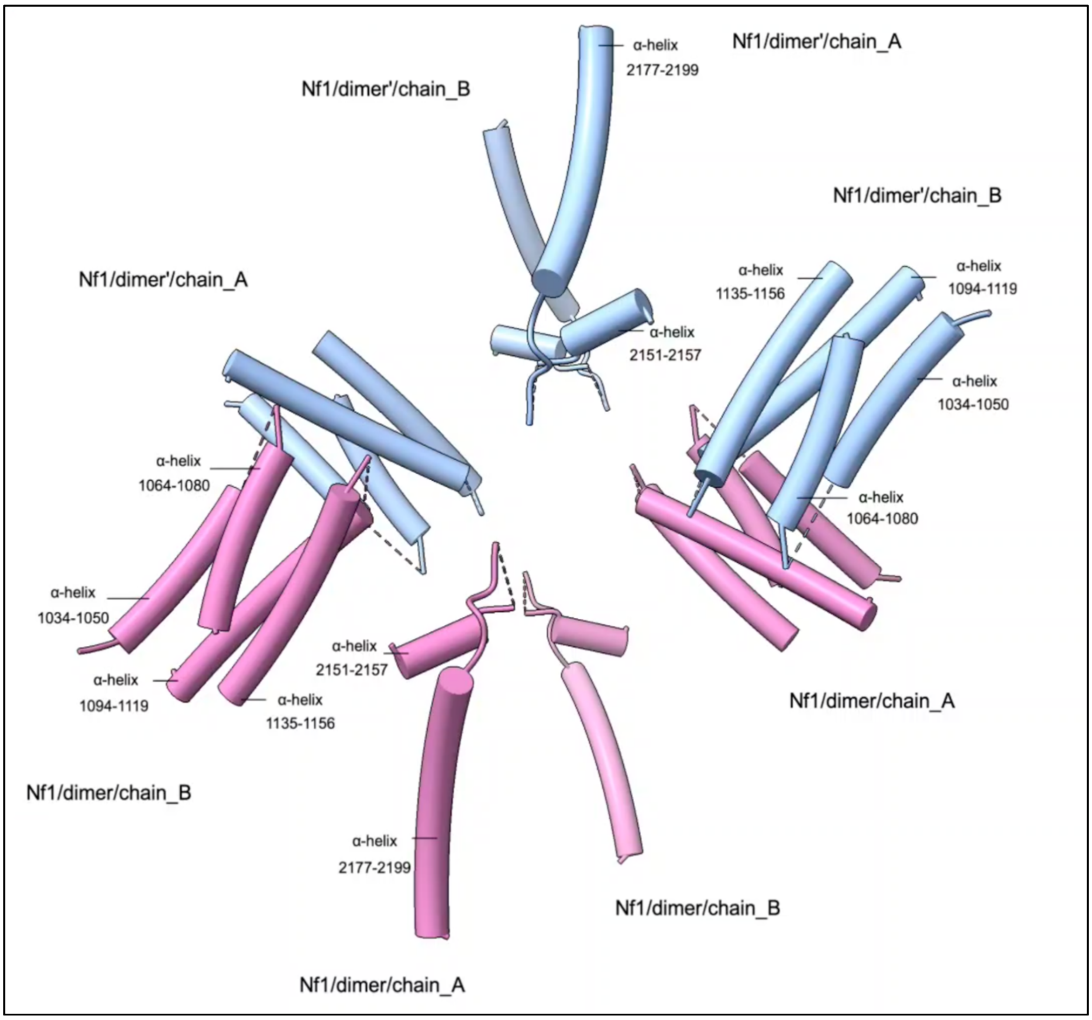


**Movie S2.** Conformation changes of the NF1 tetramer with a focus on the formed central cavity. Link to download:
https://keeper.mpdl.mpg.de/f/0536679379674fe3bee7/?dl=1
